# Optimal reproductive phenology under size-dependent cannibalism

**DOI:** 10.1101/840264

**Authors:** Nao Takashina, Øyvind Fiksen

## Abstract

Intra-cohort cannibalism is an example of a size-mediated priority effect. If early life stages cannibalize slightly smaller individuals, then parents face a trade-off between breeding at the best time for larval growth or development and predation risk from offspring born earlier. This game-theoretic situation among parents may drive adaptive reproductive phenology towards earlier breeding. However, it is not straightforward to quantify how cannibalism affects seasonal egg fitness or to distinguish emergent breeding phenology from alternative adaptive drivers. Here, we devise an age-structured game-theoretic mathematical model to find evolutionary stable breeding phenologies. We predict how size-dependent cannibalism acting on eggs, larvae, or both change emergent breeding phenology, and find that breeding under inter-cohort cannibalism occurs earlier than the optimal match to environmental conditions. We show that emergent breeding phenology patterns at the level of the population are sensitive to the ontogeny of cannibalism, i.e. which life stage is subject to cannibalism. This suggests that the nature of cannibalism among early life stages is a potential driver of the diversity of reproductive phenologies seen across taxa, and may be a contributing factor in situations where breeding occurs earlier than expected from environmental conditions.

## Introduction

The reproductive schedule of species has a tight connection to fitness and ontogenetic development [1, 2, 3], and we need to understand how size-dependent interactions and breeding phenology interacts to predict how organisms adapt to a changing climate [4]. Improving our ability to predict phenologies and seasonal structure of species interactions illuminates evolutionary traits and provides a tool for ecosystem management, conservation, and estimations of the impact of climatic change on species [4, 5, 6].

The colonization of habitats and arrival times of species or offspring in a seasonal environment is important in predator-prey interactions and in forming the structure of communities [7, 8, 9, 10, 11, 12]. For instance, a large difference in arrival times of nymphs of two dragonfly species causes the exclusion of a late arrival species [9]. This is known as priority effects, which emphasize how the sequence of breeding or other phenologies determine interaction strength. Priority effects are often size-mediated [9, 12], as organisms emerge small and grow larger while their role as predators and prey shift [13]. An early start can be a large advantage both in competitive and in predator-prey interactions, but are traded against the match with key resources during ontogeny [14].

Cannibalism is a size-mediated priority effect that occurs within species. It is ubiquitous in many groups of animals, including insects, fish, amphibians, birds, and mammals [15, 16], and can account for a major part of early-life mortality [17,18]. Intraspecific oophagy is rather common in many egg-lying animals, and eggs and newly hatched individuals are similarly vulnerable to cannibalism due to their limited ability to avoid predation and nutrient rich composition [16]. It also drives population dynamics [19, 20, 21, 22], offspring size selection [23], size distribution within the population [24], and reproductive behavior [15, 16, 25].

If there is a seasonal peak in the environmental growth or survival conditions for eggs and larvae, cannibalism during early life stages introduces a trade-off for parents between breeding at a favorable time in the season and the added predation risk caused by larger individuals born earlier. Similarly, those parents breeding earlier in the seasonal cycle may benefit by increased survival in their progeny, since they can feed on the younger and smaller offspring arriving later in the season, while availability to other resources in the environment select against earlier breeding. This mechanism may induce seemingly mal-adaptive breeding phenologies, because the cannibalistic mortality is less evident to us than the environmental drivers. As an example, a recent study pointed out that the Atlantic bluefin tuna spawn surprisingly early in the Mediterranean Sea [3] (Fig. 1). The tuna spawn at times when temperatures are lower than the optimum for growth and development for eggs and larvae. This may of course be due to other constraints, such as a food limitation, but the highly cannibalistic nature of the larvae of these tuna [26] may also contribute to this phenology. Other examples of observations where cannibalism may be shaping breeding phenology are the spatiotemporal split of larval load in the fire salamander [27], or the early hatching of damselfly [25]. Such observations suggest that cannibalism can play a major role for the optimal phenology of a wide variety of species. The frequency-dependent nature of the problem requires a non-cooperative game-theoretic approach to find the evolutionary stable strategy (ESS) [28, 29] under size-dependent cannibalism.

**Figure 1:**
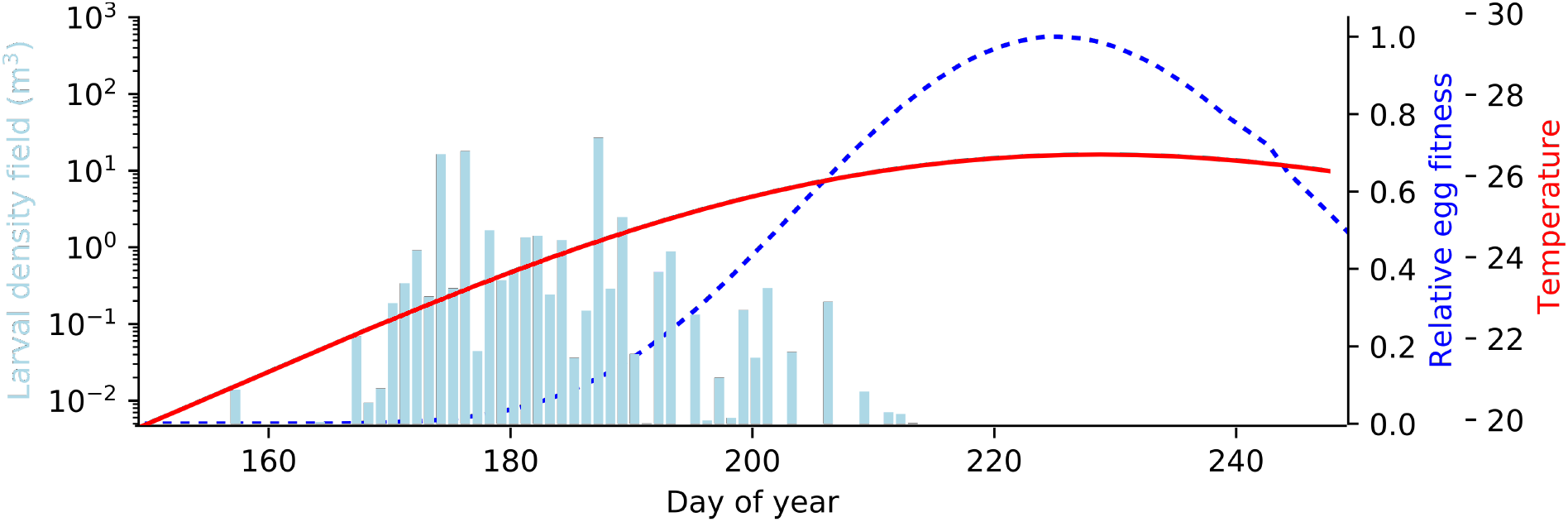
A real world example of potential cannibalistic drive towards early spawning in the highly cannibalistic Atlantic bluefin tuna larvae in the Mediterranean Sea. This shows the observed spawning phenology (larvae found in the field, bars) and the theoretical fitness of an egg (blue dashed line, scaled to the maximum value) calculated by temperature-dependent egg development time, larvae growth rate, and mortality born at any given day of the year. The red line shows the seasonal temperature cycle, which is an important driver egg and larval development rate and fitness. The figure is modified from [3].

Here we devise a theoretical approach to investigate how size-dependent cannibalism can shift breeding phenology in a seasonal environmental cycle. Our focus is on size-mediated priority effects on reproductive phenology, such as the effects of (i) intensity of cannibalism, (ii) the ontogeny of cannibalism, and (iii) duration of breeding season. We combine age-structured population dynamics with game-theoretic decision making to predict an evolutionary stable breeding phenology. Our goal is to achieve general predictions about the role of intensity and ontogeny of cannibalism, and the duration of the breeding season for breeding phenology. To keep the analysis simple, we focus only on the mortality part of cannibalism as in [19, 20, 21, 23]. This means that we do not consider resource limitation in the growth of larvae explicitly, there is no competition for food resources involved in the model, only the extra death risk involved for later born offspring. In relation to the Bluefin tuna example, this is equivalent to assuming temperature is the main environmental driver for growth and development, and that larvae will be able to find enough food even without intra-cohort cannibalism, but that they will consume smaller con-specifics at encounter. This gives a conservative assessment of the benefits of early breeding from a cannibalistic drive.

A common starting point to understand the timing of breeding or spawning in seasonal environments is the “match-mismatch” theory, where parents are selected to match their egg laying to the best environmental conditions for their offspring [30, 14]. In theoretical models, this is often inferred as a seasonal peak in food availability, which is also the best match for the critical period of the young.

We define the seasonal environment from some abiotic variable (e.g., temperature) that determine hatching success at time *t*. The optimal breeding time is to match the best environment for hatching if there is no cannibalism (Fig. 2). Then we add cannibalism among early life stages and game-theoretic decision making of breeding time among parents. If cannibalism is sufficiently strong, it can lead to deviations from the expectation of all offspring appearing at the optimal environmental conditions for hatching. The intensity of cannibalism is quantified by emerging survivorship through the egg and larval stages, depending on the number of larger individuals in the cohort. Here, we discuss three general size-dependent, ontogenetic cannibalistic modes: (i) egg are cannibalised by larvae (e.g., anuran [31] and *Tribolium* (flour beetle) [21]); (ii) larvae are cannibalised by larger larvae (e.g., damselfly [18]); and (iii) larvae cannibalise both eggs and smaller larvae (e.g., cape anchovy [32] and many fish species - see review in [33]). Understanding the potential for intra-cohort cannibalism to induce phenological shifts in the optimal breeding schedule can unveil the potential for adaptation under environmental change and explain what otherwise seem to be sub-optimal behavior.

**Figure 2:**
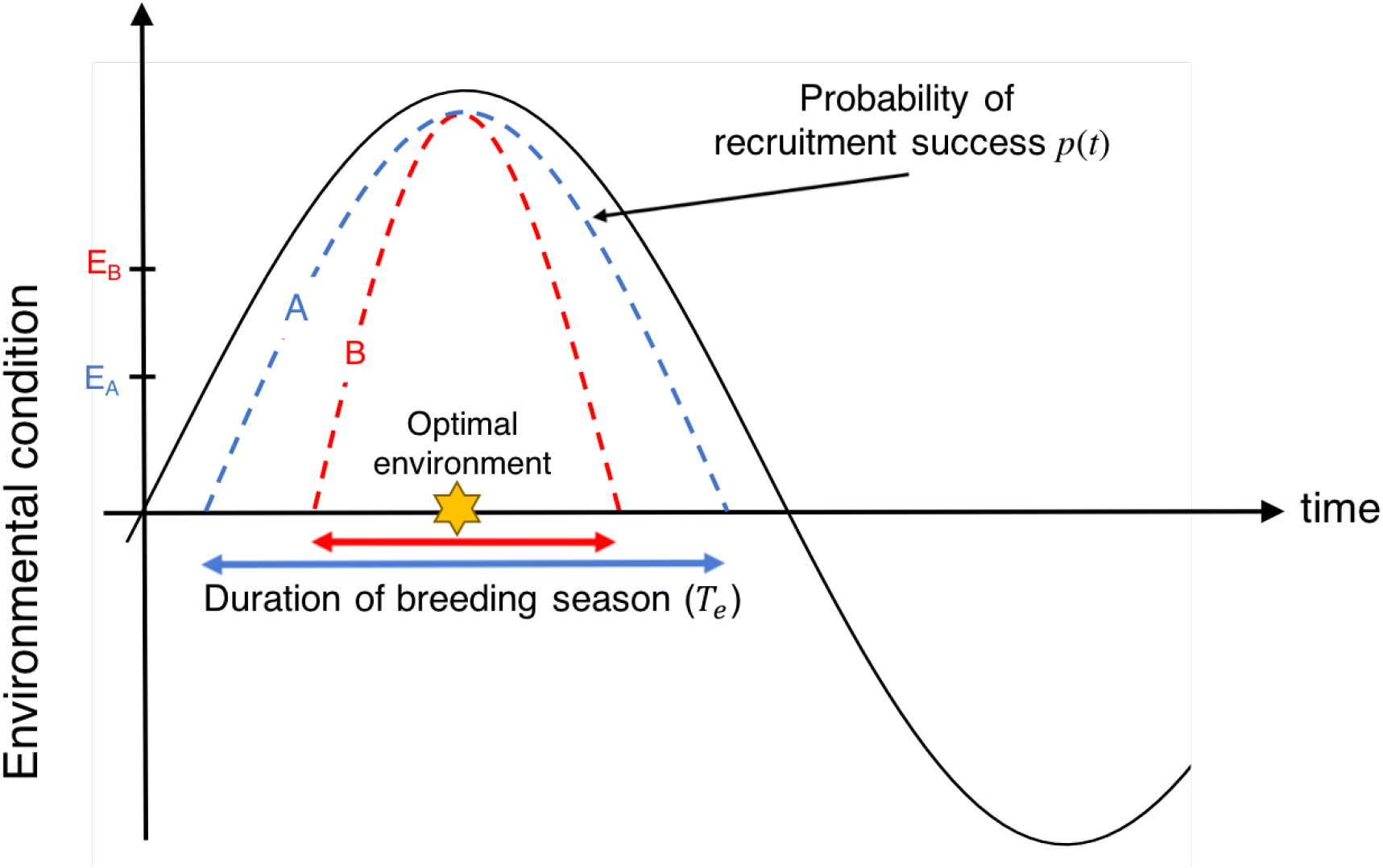
Schematic diagram of the abiotic environmental seasonal cycle and probability of recruitment success *p*(*t*) of two imaginary breeding seasons A and B where the duration of breeding seasons *T*_*e*_ are indicated by bidirectional horizontal arrows. Eggs can hatch when environmental condition is above the physiological tolerance limits in an environment (thus, egg hatching success of an egg produced at this time is larger than zero *p*(*t*) *>* 0; the physiological tolerance limits of two hypothetical breeding seasons A and B are represented by *E*_*A*_ and *E*_*B*_, respectively, on *y*-axis). The star icon represents the optimal environmental conditions for an egg to hatch, and without cannibalism we expect breeding to occur around this time to match this optimal environment with the hatching. With cannibalism, there can be a tradeoff between breeding at times of high egg hatching success or earlier, at times with lower risk of cannibalistic predation by older larvae. Therefore, there is a game between parents for optimal breeding time to maximize the survivorship of their own eggs to the adult stage.

## Material and Methods

Population dynamics that incorporate age-dependent cannibalism is often described by the McKendrick-von Foerster equation [19, 20,21]. We employ this as a basic model by assuming that older individuals are also larger, and describe the three different age-dependent cannibalistic modes: egg cannibalised by larvae, larvae cannibalised by elder larvae, and these combined. Our model describes deterministic population dynamics through early life stages and to the age at maturation *a*_*m*_. Transition to the larval stage occurs after the egg development is completed, at age *a*_*r*_. Schematic diagrams of the model are provided in Fig. 3.

**Figure 3:**
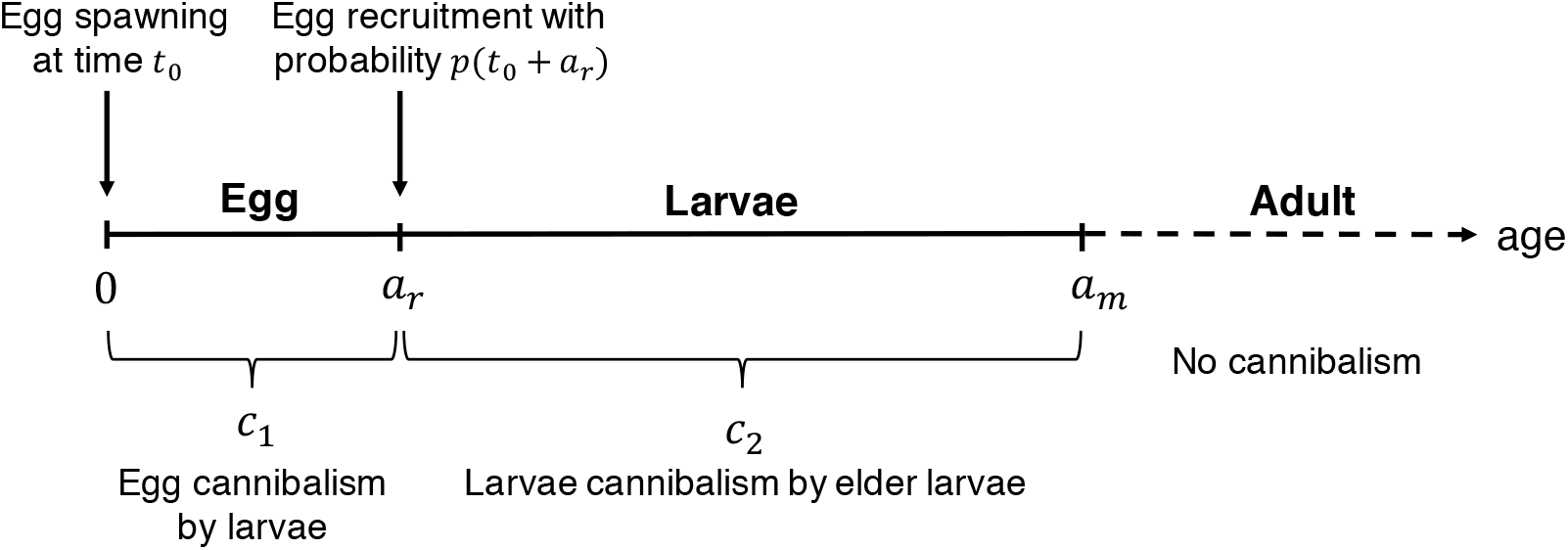
Schematic diagram of size-dependent cannibalism in the model. A constant rate of mortality from other sources is also included. We do not consider the dynamics of adult individuals and, hence, the corresponding age axis is described by the dotted line.

The dynamics of a population with age *a* at time *t* in an age-dependent cannibalistic predation model is thus described by

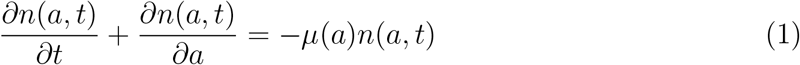

where, *μ*(*a*) is the age-specific mortality composed of the natural and cannibalistic mortalities

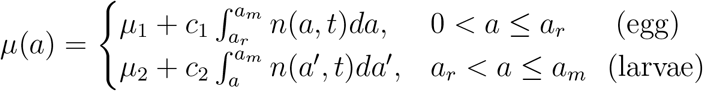

where, *μ*_1_ and *μ*_2_ are the egg and larvae natural mortality rates, respectively, *c*_1_ is the egg cannibalism rate by all larvae, and *c*_2_ is the larval cannibalism rate by elder larvae. Eq. (1) is prescribed with the boundary condition of the total number of eggs spawned at time *t*

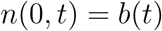

where the total number of egg *E* = ∫ *n*(0*, t*)*dt* in the breeding season is held fixed [28, 21]. This also provides an opportunity to perform the experiment with the same number of eggs [21].

The match-mismatch hypothesis normally refers to breeding phenologies adapted to a peak in some key food resource. Here, to keep the analysis simple, we let the abiotic environment drive the hatching success of an egg under the given environment at the day of birth. For example, Reglero et al. [3] observed that the probability of egg hatching in Bluefin tuna varies with temperature, and that temperature and therefore egg hatching success in their natural spawning followed a cyclic pattern over the season. The dashed line in Fig. 1 includes this and several other temperature-dependent processes, which this represents any abiotic or density-independent seasonal factor that define a breeding window in time.

We assume that the number of newly hatched eggs at time *t* is determined by the eggs spawned at time *t* − *a*_*r*_, where *a*_*r*_ is a fixed egg development time. Moreover, eggs experience the natural mortality rate *μ*_1_ during the egg development period and the egg hatching success is determined by the environmental condition (e.g., temperature or humidity) at time of hatching. Hence, the number of new larval recruits at age *a*_*r*_ is

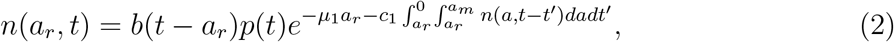

where, *p*(*t*) is the probability of hatching success at time *t*. We assume a cyclical environment as in Fig. 2, and we describe the hatching probability *p*(*t*) over the season as

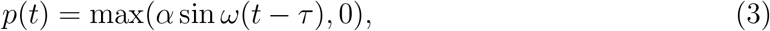

where nonnegative parameters *α* ∈ (0, 1] and *ω* determine the shape of the cycle, and *τ* is an arbitrary constant to adjust an environmental peak and the duration of the breeding season. Here, without loss of generality, we set *α* = 1. The period *T*_*p*_ defines the potential breeding period (i.e., when *p*(*t*) *>* 0).

The situation with only egg or larval cannibalism arises by setting *c*_2_ = 0 in Eq. (1) and *c*_1_ = 0 in Eq. (2), respectively. In our model, when no cannibalism occurs, the optimal time for breeding is independent of the decisions of others and it is a single point in time to match the best environment for egg hatching (Fig. 2)

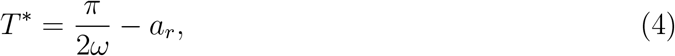

determined by the density-independent abiotic conditions.

Let us define the egg fitness at breeding time *t*_0_ as the fraction of eggs that survive to the adult stage at time *t*_0_:

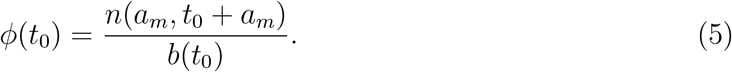

When many parents compete with each other to maximize the fitness of their offspring in a non-cooperative game-theoretic context or ESS, the equilibrium egg fitness satisfies [28]

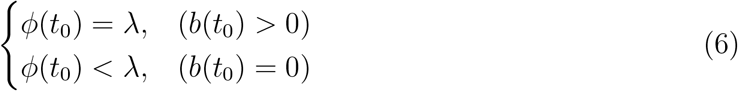

where, *λ* is a positive constant. At this equilibrium, we anticipate that egg phenology in the potential breeding season, *T*_*p*_, shows an emergent egg distribution with a length *T*_*e*_. For convenience, we define the fraction of effective breeding season as *T*_*e*_/*T*_*p*_. The deterministic population dynamics model we use (Eq. (1)) does not distinguish between single and multiple reproduction events of single individuals in one breeding season, but merely search for an egg distribution that satisfies the condition Eq. (6).

As in [28], we assume a constant number of eggs and do not consider the population dynamics of adults. The ESS breeding phenology only represents one single breeding season, since following an ESS in a dynamic population over time involve large computational challenges. To find the optimal breeding schedule under size-dependent cannibalism, we numerically integrate Eq. (1) and perform a heuristic algorithm to find the condition satisfying with Eq. (6). See Appendix for more details on the numerical details.

## Results

The intensity of cannibalism can differ between life stages, and we examined parameter sets representing larvae consuming eggs or smaller larvae or both. We also varied the length of the viable breeding season, and the total number of eggs produced by the parental population. Table 1 is the list for the parameter values examined.

**Table 1:** Parameters values used in the analysis

| Symbol | Parameter | Value |
| --- | --- | --- |
| $E$ | number of eggs | $\{10^3, 10^4, 10^5\}$ |
| $a_r$ | age at recruitment | 0.041 (15 days) |
| $a_m$ | age at maturation | 1 |
| $T_p$ | potential breeding period | $\{0.25, 0.5\}$ |
| $c_1$ | egg cannibalism rate ($\text{time}^{-1} \text{ predator}^{-1}$) | $\{0, 10^{-4}, 10^{-3}, 10^{-2}\}$ |
| $c_2$ | larvae cannibalism rate ($\text{time}^{-1} \text{ predator}^{-1}$) | $\{0, 10^{-4}, 10^{-3}, 10^{-2}\}$ |
| $\mu_1$ | egg natural mortality rate ($\text{time}^{-1}$) | 0.1 |
| $\mu_2$ | larvae natural mortality rate ($\text{time}^{-1}$) | 0.1 |

\* Results are found in Appendix.

In general, cannibalism spread the emergent egg distribution over the breeding period (i.e., spawning asymmetry), rather than at a single breeding time as in a non-cannibalistic population (Fig. 4 (a)-(c), top). Also, breeding occurs as soon as the environment allows eggs to hatch (an event that takes place after the egg development time *a*_*r*_) and most of the breeding takes place earlier than the environmental optimum (stars on *x*-axis). Hatching earlier is favored as the cannibalism intensity increases, and this trend shifts the distribution earlier in the season, leading to a narrower breeding phenology due to an accumulation of egg density towards the earliest possible breeding. This creates a monotonically decaying breeding distribution. The intensity of cannibalism is also characterized by the egg fitness to the age at maturation, *a*_*m*_ (Fig. 4 (a)-(c), bottom). Cannibalism on eggs causes lower egg fitness, and earlier hatching is favored, and more than for larval cannibalism. The emerging breeding period also varies with the strength of cannibalism, and it is more contracted with higher cannibalism.

**Figure 4:**
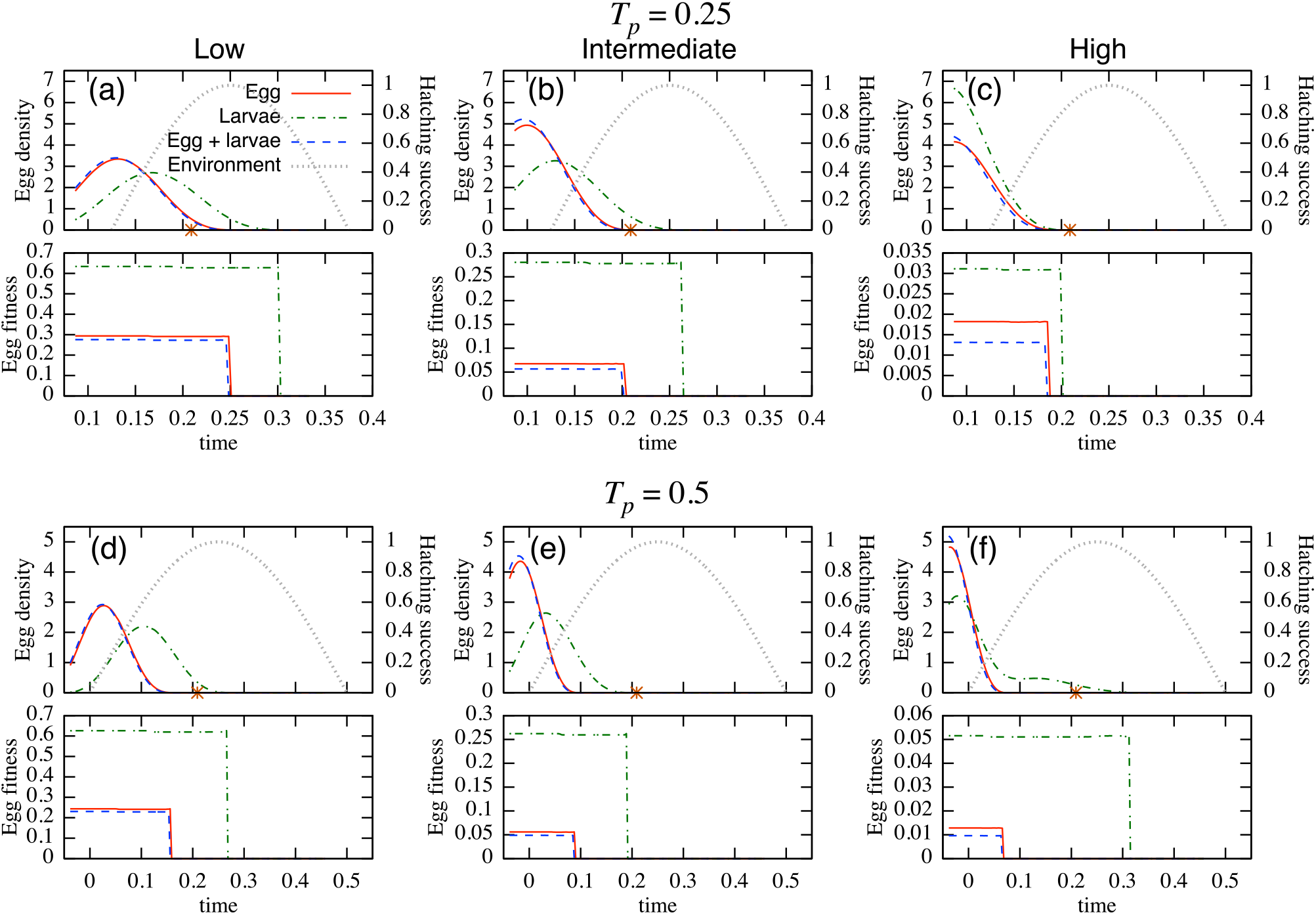
The distributions of the normalized egg density at breeding time (top of each panel: left-vertical axis) under three cannibalism target (i) egg cannibalised by larvae (Egg); (ii) larvae cannibalised by larger larvae (Larvae); and (iii) larvae cannibalise both eggs and smaller larvae (Egg+larvae), and egg recruitment success in relation to the environmental condition (right-vertical axis). Bottom of each panel is egg fitness. These are examined under three cannibalistic scenarios: low (left; *c*_1_ = *c*_2_ = 0.0001); intermediate (center; *c*_1_ = *c*_2_ = 0.001); and high (right; *c*_1_ = *c*_2_ = 0.01). The stars on the *x* axis on the top of each panel represent the optimal spawning time under the no cannibalistic predation. The potential breeding period is *T*_*p*_ = 0.25 (top two rows) and *T*_*p*_ = 0.5 (bottom two rows). The number of eggs is *E* = 10^4^. Other parameter values are shown in Table 1.

We observed similar trends when the environment suitable for breeding is increased to 50% of a year (Fig. 4 (d)-(f)). Counterintuitively, the longer potential hatching periods tend to result in shorter breeding periods. An exception is when cannibalism is high and only on larvae (Fig. 4 (f)). In this case, the distribution of the breeding period is wider when the environmentally suitable breeding window is longer, giving a bimodal breeding distribution with one peak close to the abiotic environmental optimum and one peak early in the season, a signal of a breeding cycle within a single breeding period. Average values and fractions of effective breeding seasons are provided in Table 2 and the qualitative discussions above are verified.

**Table 2:** Mean value and the fraction of effective breeding season, *T*_*e*_/*T*_*p*_, of each curve in Figures 4 (mean, *T*_*e*_/*T*_*p*_).

|  | egg | larvae | egg & larvae |
| --- | --- | --- | --- |
| $T_p = 0.25$ | | | |
| Low | (0.141, 0.659) | (0.171, 0.868) | (0.139, 0.648) |
| Intermediate | (0.120, 0.473) | (0.142, 0.714) | (0.118, 0.462) |
| High | (0.0743, 0.407) | (0.114, 0.462) | (0.069, 0.396) |
| $T_p = 0.5$ | | | |
| Low | (0.0324, 0.396) | (0.106, 0.615) | (0.030, 0.390) |
| Intermediate | (-0.00, 0.258) | (0.0412, 0.462) | (-0.002, 0.253) |
| High | (-0.008, 0.214) | (0.0364, 0.709) | (-0.009, 0.209) |

Qualitatively similar patterns appear with different total numbers of eggs, *E* (see Figs. A.1 for *E* = 10^3^ and A.2 for *E* = 10^5^, respectively, in Appendix). The exception is again when the cannibalistic intensity is high, and it is only imposed on larvae, and if the total number of eggs is ten times larger (*N* = 10^5^; Figs. A.2 (c) and (f)). Then, the smaller egg fitness yields a wider emergent breeding period.

## Discussion

Cannibalism on egg, larvae, or both from larvae hatched earlier in the season lead to intraspecific competition over the optimal breeding period, and apparently suboptimal breeding phenologies relative to some abiotic optimum may emerge. Firstly, cannibalism causes a breeding asymmetry rather than a simple match to the best seasonal environment. Emergent breeding phenologies are the outcome of a game-theoretic intraspecific competition between parents where each individual make a decision based on decisions of the others. It differs from other mechanisms inducing hatching asymmetry, such as conservative bet-hedging [34] and diversified bet-hedging strategies [35] where the main goal is adaptation to variable environments, not involving the choice of con-specifics [36]. Secondly, the model predicts a shift of breeding towards earlier and less favourable conditions for the progeny if cannibalism is an important source of mortality. These two properties agree with observations of the spawning phenology of Atlantic bluefin tuna in the Mediterranean Sea [3], where size-dependent larval intra-cohort cannibalism is known to occur [26]. Also, DeBlock et al. [25] observed early and log-normally distributed breeding phenologies under egg cannibalism in damselfly populations.

Emergent breeding phenology is sensitive to the ontogeny of cannibalism. Our model predicted that egg cannibalism tends to induce an earlier shift than larval cannibalism, since later hatching causes greater cannibalistic risk by the larvae. Hence, if eggs are vulnerable to intra-cohort cannibalism then total predation from cannibals are higher and the drive towards earlier spawning is stronger. For cannibalism acting on larvae, on the other hand, the cannibals are themselves thinned out by cannibalism. This limits the magnitude of cannibalistic mortality among smaller larvae, and allows parents to breed later and under a better environmental conditions.

A longer potential breeding period (*T*_*p*_ = 0.5) gave a lower ratio of emergent:potential breeding season (*T*_*e*_/*T*_*p*_) than the shorter potential breeding period (*T*_*p*_ = 0.25). In a shorter potential breeding period, the tradeoff between the better seasonal environment and reduced predation risk is weaker, and provides a higher benefit to individuals breeding later in the season than the case of a longer potential breeding period, leading to less contracted breeding phenologies. The magnitude of change in the ratio again depends on cannibalism rate.

Also, depending on the intensity of cannibalism over the ontogeny, cannibalism can induce surprisingly diverse patterns of emergent breeding phenologies, including a symmetric bell-shaped distribution with and without cut off at the earliest possible breeding time or monotonically decreasing egg production curves (e.g., Fig. 4 (a) and (c), top). The latter distribution appears in response to intensified cannibalism, and as a rule of thumb, intensifying cannibalism causes a shift of the egg distribution toward an earlier time and it leads to a cut-off of the distribution at the earliest possible breeding time (e.g., Fig. 4 (d)-(f), top). Moreover, even a bimodal pattern, a signal of a breeding cycle within a single breeding period (e.g., Fig. 4 (f), top) or more complex phenologies (Figs. A.2 (c) and (f), top) can occur if cannibalism is only on larvae by elder individuals and its effect on survival is high. These diverse patterns were previously reported in multiple taxa, and may be attributed to highly nonlinear mechanisms in reproductive decision making. The observed spawning phenology of Atlantic bluefin tuna resembles a bell-shaped curve [3], while a monotonically decreasing pattern was observed in the burying beetle population under the filial cannibalism [37]. In the context of an optimal breeding schedule of male butterflies to facilitate mating success, Iwasa et al. [28] predicted a bell-shaped pattern with a cut-off as the optimal schedule.

We have used a deterministic model to analyse the effect of cannibalism on breeding phenology. Provided with a good understanding of the lifecycle and the cannibalistic interactions of concerned species, a wide variety of tactical models can be constructed based on our general model (e.g., [38]) such as size/time dependency in biological parameters, or a more mechanistic formulation of the process of cannibalism. For example, Reglero et al. [3] experimentally showed that temperature affects the egg hatching success of the Atlantic bluefin tuna, but also egg developmental time, larvae growth and mortality rate. The tuna spawn earlier than expected from the abiotic environment, and they proposed intra-specific cannibalism as a possible cause.

While our results are based on the reproductive phenology in a single season under a deterministic environment, this can still serve general understanding of the phenology over multiple years if the environmental fluctuation is moderate, and the population dynamics is stable. However, ideally the critical period concept of the match-mismatch theory should include dynamics of the full life-cycle over multiple generations (e.g. [39]). In addition, a variable environment can lead to bet-hedging reproductive strategies [36, 35], and under such conditions, the realized phenology may deviate from our deterministic predictions. To assess these factors, we need models of greater complexity, and to connect game-theoretic and bet-hedging concepts.

Our minimal approach, assuming cannibalism causes only an additional mortality, predicts an early-shift in breeding phenology. It does not consider the benefit of energy gain by cannibalism, which may further promote an even earlier shift of breeding phenology. A physiologically structured population model is one way to explicitly include the energy gain from cannibalism [24], although the model requires greater model complexity and larger number of parameters.

Our model can also help us understand context-dependent trajectories of species assemblage and priority effects [40, 10]. Species emerge in seasons in a constant game with other species, and the interactions are size- and time-dependent. Extensions of our modelling approach can include size-dependent competitive interactions between several species, and thus become a tool to integrate evolutionary dynamics of phenologies into analysis of environmental change. Phenological response to climate change differs in a complex manner between species and it can lead to a trophic mismatch [41, 42]. Yang and Rudolf [4] emphasized the importance of understanding the interaction between phenology and stage-structured species interaction to predict the response to climatic change. Our model is only a starting point, but demonstrates the role that size-dependent intraspecific interactions can have in forming life histories and breeding phenology.

## Acknowledgements

We thank M. H. Cortez and T. Fung for reading a draft and providing thoughtful comments. We also thank two anonymous reviewers for their constructive comments.

## Data Accessibility

The manuscript does not contain original data. Computer code written in C is available upon request.

## Appendix Heuristic optimization

Here we present the numerical algorithm that heuristically searches the distribution of egg production that satisfy the condition in Eq. (6). The algorithm is based on the following simple steps

1. Prepare an arbitrary initial egg distribution where the total eggs are *E* over the potential breeding period *b*(*t*_*i*_)Δ*t* = *n_t_i__*, where *b*(*t*_*i*_)Δ*t* is the number of eggs between the time *t*_*i*_ and *t*_*i*_ + Δ*t* and *b*(*t*)*dt* = *E* holds.
2. Perform a numerical integration of Eq. (1) with the initial egg distribution and calculate fitness at each breeding schedule *ϕ*(*t*).
3. Find the maximum *ϕ*_max_ and minimum *ϕ*_min_ fitnesses among the possible breeding schedules and replace the number of eggs *dE* from the time with the lowest fitness to the highest fitness.
4. Return to step 2. until the relative difference between *ϕ*_max_ and *ϕ*_min_ becomes smaller than the predetermine threshold 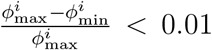, where the index *i* represents the number of iterations of this algorithm.

To avoid a cyclic egg replacement between two or more breeding times, we shuffle the egg number *dE* between two randomly chosen breeding times for every 10000 iterations. Moreover, we calculate the relative difference of the average relative fitness difference defined in step 4 for every 300 iterations over 100 iterations, and decrement *dE* by 5% when this relative difference becomes smaller than the predetermined threshold 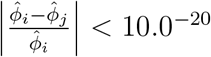, where 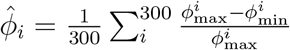 is the average relative fitness difference where its sampling starts at iteration time *i*.

Values used in this algorithm (e.g., predetermined thresholds) are heuristically determined. We used a uniform initial egg distribution. We observed the order of iterations to obtain convergence spans 10^4^−10^6^ depending on the cannibalistic parameter used in the main text. We choose this algorithm because we observe that our objective function 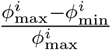 largely fluctuates over iterations (i.e., not monotonically decreases with *i*) and, therefore, ordinary optimization algorithms can not be directly applied. It is also noted that we truncated the fitness value where an egg load is smaller than 0.5, and define the interval of population *n*_*t*_(*a, t*) = max(*n*_*t*_(*a, t*), 0) to facilitate the convergence of our heuristic algorithm.

**Figure A.1:**
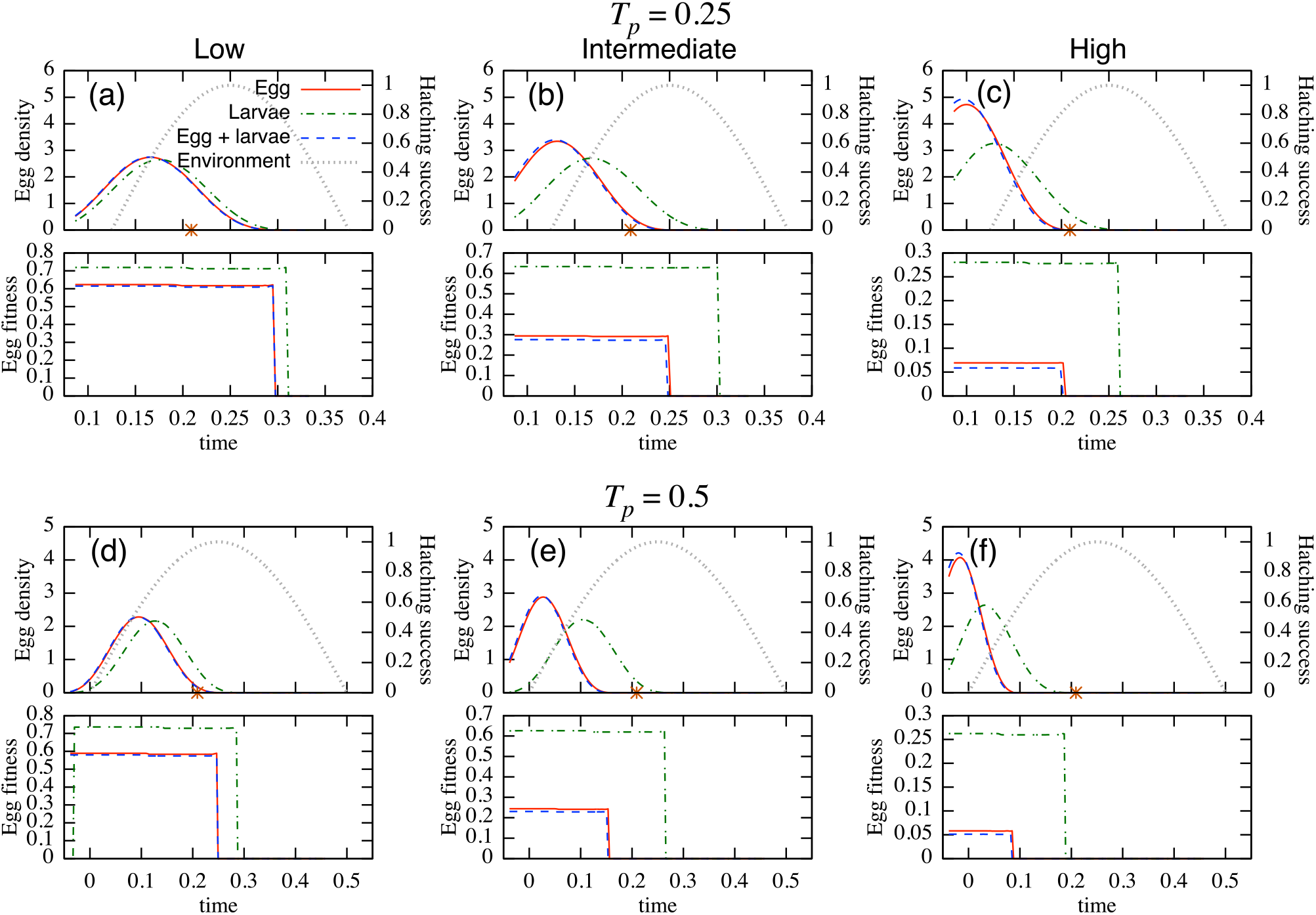
The distributions of the normalized egg density (top of each panel: left-vertical axis) under three cannibalism target (i) egg cannibalised by larvae (Egg); (ii) larvae cannibalised by larger larvae (Larvae); and (iii) larvae cannibalise both eggs and smaller larvae (Egg+larvae), and egg recruitment success in relation to the environmental condition (right-vertical axis). Bottom of each panel is egg fitness. These are examined under three cannibalistic scenarios: low (left; *c*_1_ = *c*_2_ = 0.0001); intermediate (center; *c*_1_ = *c*_2_ = 0.001); and high (right; *c*_1_ = *c*_2_ = 0.01). The stars on the *x* axis on the top of each panel represent the optimal spawning time under the no cannibalistic predation. The potential breeding period is *T*_*p*_ = 0.25 (top two rows) and *T*_*p*_ = 0.5 (bottom two rows). The number of eggs is *E* = 10^3^. Other parameter values are shown in Table 1.

**Figure A.2:**
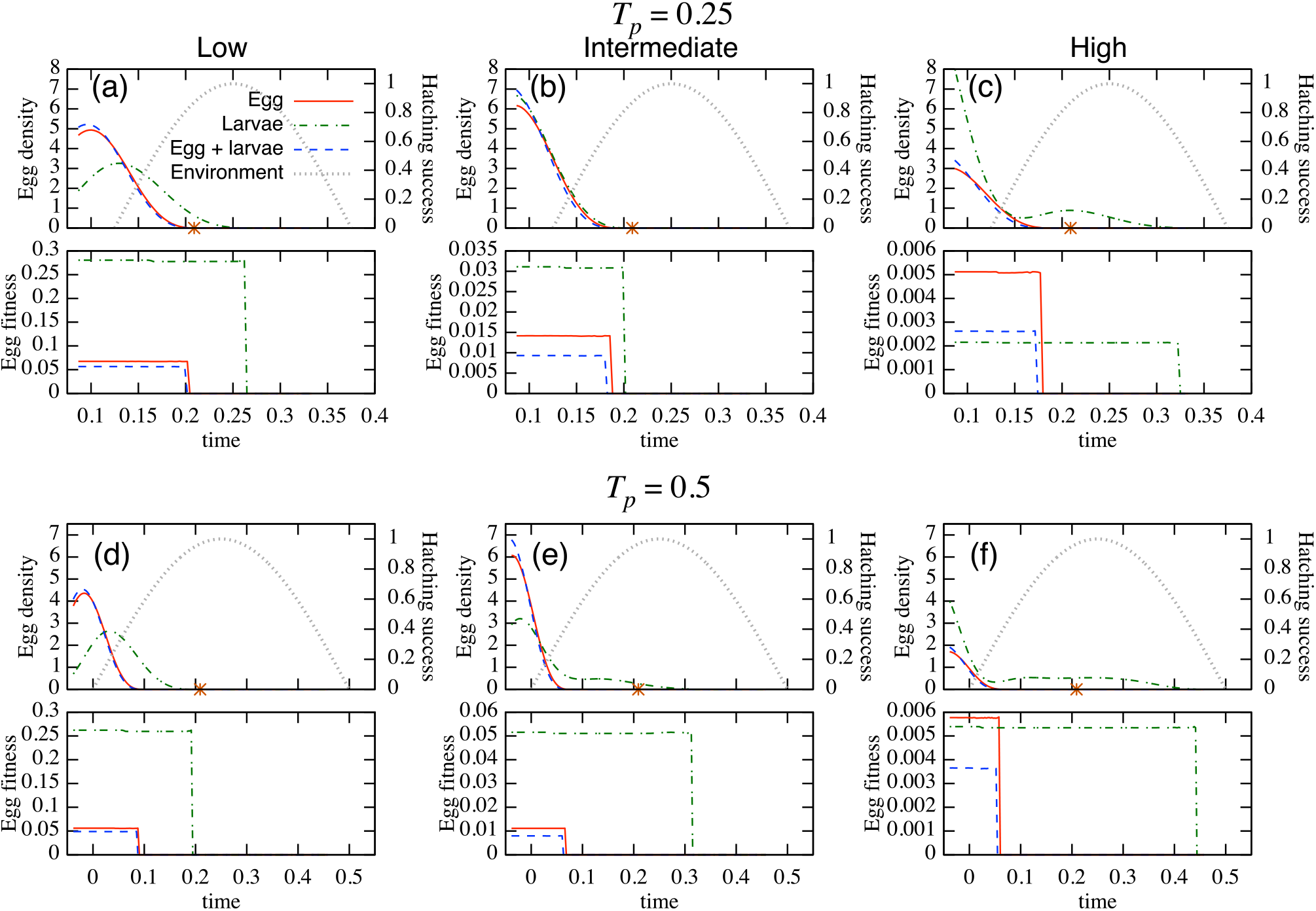
The distributions of the normalized egg density (top of each panel: left-vertical axis) under three cannibalism target (i) egg cannibalised by larvae (Egg); (ii) larvae cannibalised by larger larvae (Larvae); and (iii) larvae cannibalise both eggs and smaller larvae (Egg+larvae), and egg recruitment success in relation to the environmental condition (right-vertical axis). Bottom of each panel is egg fitness. These are examined under three cannibalistic scenarios: low (left; *c*_1_ = *c*_2_ = 0.0001); intermediate (center; *c*_1_ = *c*_2_ = 0.001); and high (right; *c*_1_ = *c*_2_ = 0.01). The stars on the *x* axis on the top of each panel represent the optimal spawning time under the no cannibalistic predation. The potential breeding period is *T*_*p*_ = 0.25 (top two rows) and *T*_*p*_ = 0.5 (bottom two rows). The number of eggs is *E* = 10^5^. Other parameter values are shown in Table 1.

**Table A.1:** Mean value and the fraction of effective breeding season, *T*_*e*_/*T*_*p*_, of each curve in Figures A.1 (mean, *T*_*e*_/*T*_*p*_).

|  | egg | larvae | egg & larvae |
| --- | --- | --- | --- |
| $T_p = 0.25$ | | | |
| Low | (0.169, 0.846) | (0.178, 0.901) | (0.168, 0.846) |
| Intermediate | (0.141, 0.659) | (0.171, 0.868) | (0.139, 0.648) |
| High | (0.116, 0.473) | (0.142, 0.703) | (0.113, 0.462) |
| $T_p = 0.5$ | | | |
| Low | (0.096, 0.577) | (0.126, 0.637) | (0.095, 0.577) |
| Intermediate | (0.032, 0.390) | (0.106, 0.610) | (0.030, 0.385) |
| High | (0.000, 0.253) | (0.041, 0.456) | (-0.002, 0.247) |

**Table A.2:** Mean value and the fraction of effective breeding season, *T*_*e*_/*T*_*p*_, of each curve in Figures A.2 (mean, *T*_*e*_/*T*_*p*_).

|  | egg | larvae | egg & larvae |
| --- | --- | --- | --- |
| $T_p = 0.25$ | | | |
| Low | (0.120, 0.473) | (0.142, 0.747) | (0.118, 0.462) |
| Intermediate | (0.106, 0.407) | (0.114, 0.462) | (0.102, 0.385) |
| High | (0.045, 0.374) | (0.136, 0.956) | (0.041, 0.352) |
| $T_p = 0.5$ | | | |
| Low | (-0.001, 0.258) | (0.041, 0.467) | (-0.002, 0.253) |
| Intermediate | (-0.010, 0.214) | (0.036, 0.709) | (-0.012, 0.203) |
| High | (-0.003, 0.198) | (0.101, 0.967) | (-0.004, 0.187) |

